# Alpha-2 agonism of the locus coeruleus impairs learning driven by negative prediction error

**DOI:** 10.1101/2024.09.13.612879

**Authors:** Simon K. C. Lui, Ashleigh K. Brink, Laura H. Corbit

**Author notes:** **Corresponding Author:** Laura Corbit, Department of Psychology, Department of Cell & Systems Biology, University of Toronto, Toronto, Ontario, Canada, M5S 3G5.

## Abstract

Refining previous learning when environmental contingencies change is a critical adaptive function. Studies have shown that systemic noradrenaline (NA) manipulations, as well as optogenetic manipulations of the locus coeruleus (LC), the primary source of forebrain NA, can strengthen long-term retention of appetitive extinction. To determine whether the contribution of NA is specific to extinction or extends to other forms of learning where reward is less than expected, we suppressed LC activity with clonidine, an α2A-adrenergic receptor agonist, in two tasks: compound extinction, where two previously rewarded cues are paired and no longer rewarded, and overexpectation, where animals are presented with two previously rewarded cues but receive a single reward rather than the expected two. In compound extinction, we found no differences between groups in training, extinction, or a spontaneous recovery test. However, animals that received clonidine reacquired responding to the previously extinguished cue significantly faster than saline animals, suggesting weakened extinction learning. In overexpectation testing, the saline group responded significantly less to a stimulus that had undergone overexpectation relative to a control stimulus, indicating that they had recalibrated their estimation of reward magnitude following training where reward was less than expected. In contrast, clonidine-treated animals did not differ in responding to the overexpectation versus control stimuli, suggesting that clonidine impaired learning resulting from overexpectation. These results demonstrate that activity of the LC is important for learning to reduce responding in both extinction and overexpectation paradigms.

---

Detecting and encoding relationships between events as well as modifying previous learning when confronted with new information is essential for adaptive control of behaviour. For example, when a previously reinforced stimulus or response is no longer reinforced, the previously learned behaviour should be reduced or extinguished, allowing resources to be allocated elsewhere. However, there is substantial evidence that the expression of extinction is often not permanent, and the original learning can return [1–3]. When the originally learned behaviours are unwanted, or no longer appropriate, this presents a challenge for behaviour modification, particularly in the clinical context of extinction-based therapies. Thus, there is interest in strategies to enhance the long-term expression of extinction.

Consideration of the factors that cause learning in the first place can identify strategies for improving learning. Contemporary theories suggest that learning is driven by the discrepancy between what is expected based on available predictors, and what actually occurs[4–6]. This discrepancy is often referred to as prediction error and the larger the error, the more learning that should result on a given trial. Two paradigms exploit this idea; compound extinction and overexpectation. In either task, multiple stimuli are trained independently, and each comes to predict the reinforcer it has been paired with. When such stimuli are then presented simultaneously, each predicts the outcome and so together, double the outcomes are predicted. When this expectation is not met, either in extinction when no reinforcer is delivered, or in overexpectation, when the original, but smaller than expected reinforcer is delivered, a negative prediction error is produced leading to a decrease in the associative strength, or predictive power, of each stimulus across trials[5,7–10].

Extinction and overexpectation provide methods for reducing previously learned behaviours, however, both are subject to spontaneous recovery and renewal [3,11–13], suggesting that their expression can be disrupted by the passage of time or changes in the test environment, and that they rely on new inhibitory learning that competes for behavioural control, rather than the erasure of the original learning. However, compound extinction has been shown to be more resistant to spontaneous recovery than extinction of a single stimulus [14–17], and augmenting noradrenergic activity has been found to enhance appetitive extinction [15–17], while inhibition of NA signalling with propranolol, a beta-adrenergic antagonist, can impair extinction of rewarding stimuli [15–18]. Thus, these strategies have potential for producing more lasting behavioural change. Nonetheless, the neural and pharmacological bases of these effects are not well understood.

Noradrenaline (NA) has been shown to contribute to the formation of strong memories for emotionally salient events [19,20]. Microdialysis investigations have observed an increase in extracellular NA during extinction training [21] and voltammetry experiments have reported release of NA when an expected reward is omitted [22]. However, these studies have yet to isolate the source of these noradrenergic effects. The primary candidate is the locus coeruleus (LC), a bilateral pontine nucleus which sends noradrenergic projections throughout the forebrain [23–25], facilitating plasticity in diverse projection regions [24,26–29]. Electrophysiological and fiber photometry recording studies have observed that LC neurons resume firing in early extinction trials [30–32]. However, the literature remains mixed, as other studies have not detected this firing [33] and previous lesion studies have also produced mixed results [34–36]. Recent studies using chemo-or optogenetic LC stimulation are similarly conflicting [29,37–39], perhaps due to differences in the effects of tonic vs. phasic firing [25,40], previously unrecognised functional heterogeneity amongst LC projections [24,29], or differences in the intensity of LC stimulation and interactions with arousal state and corresponding levels of endogenous NA. Extended low frequency, or tonic stimulation, of the LC can be anxiogenic [41] and high levels of stress and corresponding NA are known to interfere with extinction of conditioned fear [37,38]. In contrast, brief higher frequency, or phasic LC stimulation, can facilitate learning and plasticity in downstream regions [42–44]. Notably, our recent work [39] found that phasic stimulation of the LC was sufficient to enhance extinction of food seeking. Thus, different levels of LC activation may produce different effects on extinction learning.

The current study sought to better understand the role of the LC in extinction learning. To do so, we infused clonidine, an alpha-2-receptor agonist into the LC to inhibit LC-NA neurons. We hypothesised that clonidine acting on local autoreceptors during extinction of food seeking would attenuate extinction learning. Furthermore, as emerging evidence now suggests that the LC plays a critical role in prediction error signaling [26,27,32,45,46] – that is, signalling that occurs when there is a difference between the expected and experienced outcome [5], we also examined whether the LC contributes to another example of learning driven by a negative prediction error; overexpectation, where reinforcement is still delivered, but is smaller than anticipated based on the stimuli presented. We infused clonidine prior to overexpectation trials, hypothesising that attenuating LC-NA activity would impair the animals’ ability to update their previous learning in this task.

## Materials & Methods

### Subjects

Forty-five (41 male, 4 female) 8-week-old Long-Evans rats served as subjects (Charles River, St. Constant, QC, Canada). Equal numbers of males and females were included in an initial cohort but because of excessive bleeding in females, males were used thereafter. Housing was identical to our previous studies [39]. All experimental procedures were performed in accordance with guidelines from the Canadian Council on Animal Care and approved by the University of Toronto Animal Care Committee.

### Apparatus

Rats were trained and tested in 8 identical behavioural chambers (Med Associates, Fairfax, VT, USA) housed within sound- and light-attenuating shells. Each chamber contained two key lights situated above retractable levers and a recessed magazine located in the center of the same wall where 45-mg dustless precision grain pellets could be delivered (F0165; BioServ, Flemington, NJ, USA). The chambers were equipped with a white noise generator and solenoid that delivered a 5-Hz clicker stimulus that were used as auditory stimuli in addition to the visual key lights. The auditory stimuli were adjusted to 80 dB in the presence of background noise of 60 dB provided by a ventilation fan. Each chamber was illuminated by a 3-W, 24-V house light. All experimental events including stimulus delivery were controlled with MED-PC V software (Med Associates), which also recorded magazine entries and lever-press responses.

### Stereotaxic Surgery

General surgical procedures have been described elsewhere [39]. Briefly, stereotaxic surgery was performed under aseptic conditions to introduce guide cannulae to bilaterally target the LC (AP: -10.0 mm, ML: ±1.3 mm, DV -7.4 mm, relative to bregma and skull). Rats were anesthetized with isoflurane (5% induction, 2-3% maintenance).

Meloxicam (2 mg/kg) was administered subcutaneously prior to surgery for analgesia. In an attempt to reduce bleeding, a subset (n = 5) of animals had the LC targeted at a 20º angle posterior to vertical and coordinates adjusted to ensure similar placement within the LC (AP: -12.90 mm, ML: ±1.3 mm, DV: -7.77 mm, relative to bregma and skull). A bilateral guide cannula (26G; HRS Scientific, Anjou, QC, Canada) was lowered to a position 3.0 mm above the LC and secured in place with dental cement. The internal cannula used for drug delivery extended 3.0 mm past the guide cannula to reach the final DV coordinates. Dummy cannulae were inserted into the guides to prevent blockage. A single dose of dexamethasone (1 mg/kg) was given immediately after surgery. Additional meloxicam (2 mg/kg) was administered 24 and 48 hours postoperatively. Rats were given a minimum of 1 week for recovery before commencement of any behavioural training.

### Behavioural Training & Testing

All rats were first trained in a discriminated operant task where lever-pressing in the presence of a discriminative stimulus resulted in food reward. Clonidine was delivered prior to key extinction or overexpectation sessions as outlined below.

#### Magazine & Lever Training

Following surgical recovery, rats were food restricted and maintained at 90% of their free feeding weight. On the first training day, rats received a 30-minute magazine training session where approximately 30 pellets were delivered according to a random time 60-second schedule (RT-60s). The following day, rats were trained to lever-press for food reward. Each lever press delivered a pellet until 50 pellets were earned, at which point the session was terminated. Rats that failed to earn 50 pellets were given a second identical session the following day.

#### Discrimination Training

Next, rats were trained in a discriminated operant procedure in which responding during stimulus presentations resulted in pellet delivery, whereas responding in the absence of a stimulus had no programmed consequences. Each session contained eight, 20-second presentations of a light (both key lights illuminated), white noise, or clicker stimulus (24 trials total). To aid acquisition, on the first day the lever extended during stimulus presentations and retracted when the stimulus ended. On subsequent days, the lever was present throughout the entire session but only responding during the stimuli was reinforced. All rats were trained on random ratio (RR) schedules that increased across days (3 days of continuous reinforcement, 3 days of RR2, and 3 days of RR4). Trial order was pseudo-randomly determined and the intertrial interval (ITI) averaged 90 seconds. Rats received two mock infusions prior to the first day of drug delivery where rats were gently held, and an internal cannula was lowered into the injection site but no infusion was given.

#### Drug Infusions

On infusion days, animals received a bilateral infusion of saline or clonidine (0.6 μg dissolved in 0.2 μL saline per hemisphere; Sigma-Aldrich, Oakville, ON, Canada) into the LC via internal cannulae (33G; HRS Scientific). This dose was selected based on pilot experiments (see Supplemental Materials, Figure S1). A total volume of 0.2 μL was infused over the span of 2 minutes, and the internal cannulae remained in place for an additional 1 minute after the infusion was completed to allow diffusion away from the tip. To minimize variation related to timing of drug delivery, on drug days animals were trained and tested in groups of 4, with the delay between drug infusion and testing ranging from 5 to 20 minutes.

##### Experiment 1: Compound Extinction

Based on our previous pharmacological studies implicating noradrenaline in the ability of compound extinction to enhance subsequent expression of extinction learning [15], we examined whether LC inhibition would impair this effect. The experimental design is summarized in Figure 1. Following discrimination training, rats (n = 24) were given 2 days of extinction. These sessions were identical to the training sessions, except that lever presses were not reinforced. Prior to a third extinction session, rats were assigned to saline (n = 12) or clonidine (n = 12) groups matched for response rates on the final day of training. Following infusions, both groups received six compound stimulus presentations consisting of the light stimulus coincident with either the white noise or the clicker stimulus (counterbalanced in an effort to match groups based on response rates in training). Four weeks later rats were tested for spontaneous recovery. The test consisted of six presentations of the auditory stimulus that was presented in the final extinction session as part of the compound, but delivered alone at test (i.e., white noise or clicker). Based on the previous observation that an extinguished response can recover rapidly when the response is reinforced again [47], providing additional evidence that extinction does not erase the original learning, the following day, rats were tested for reacquisition of responding where lever-pressing during stimulus presentations was again reinforced according to a RR4 schedule.

**Figure 1.**
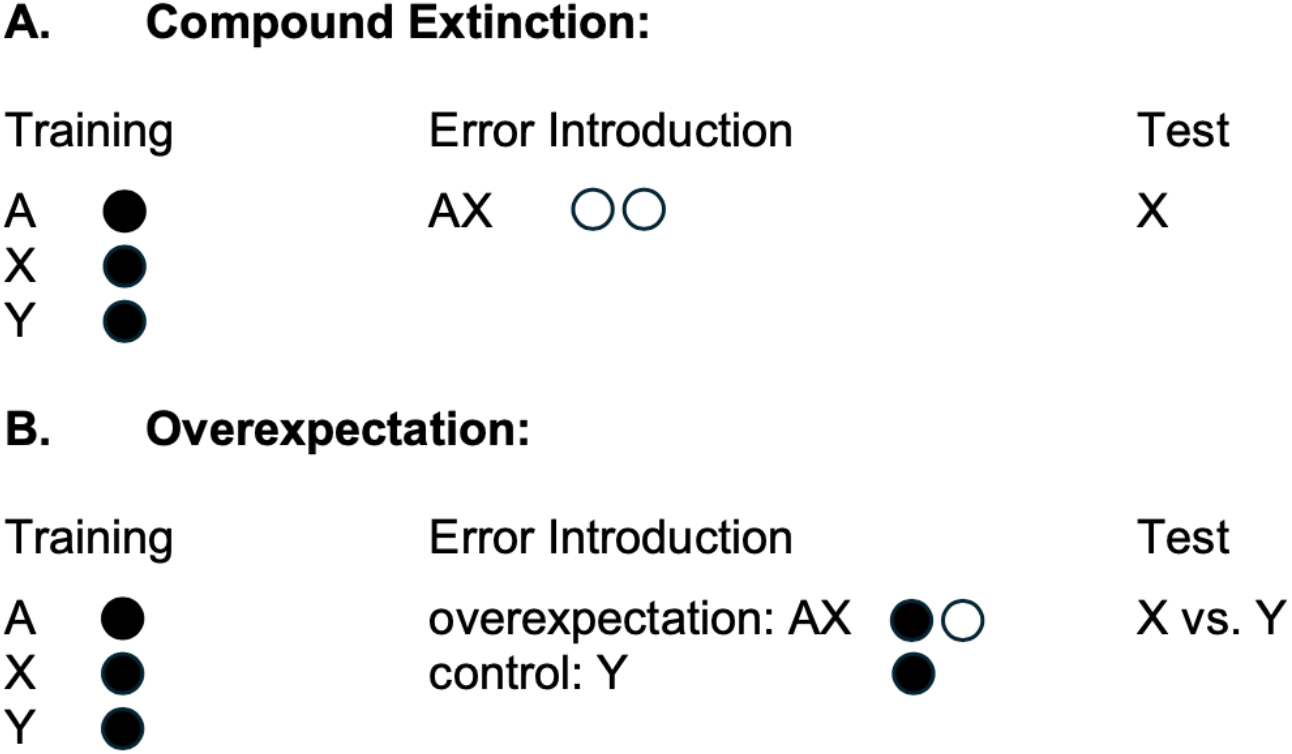
Summary of Behavioural Designs. **(A)** Training and testing procedures used in compound extinction. **(B)** Training and test procedures used in overexpectation. A respresnts the light stimulus (two key illuminated on the chamber wall.) X and Y represent the two auditory stimuli, clicker and white noise, counterbalanced. Each filled circle represents delivery of a single pellet whereas open circles represent omission of expected pellets. Saline or clonidine was infused prior to the Error introduction phase in each experiment. Saline and Clonidine groups were tested drug-free and responding to stimulus X (compound extinction) or X and Y (overexpectation) was compared.

##### Experiment 2: Overexpectation

Next, we tested the effects of LC inactivation on overexpectation in a separate cohort of rats (n = 21). In addition to the initial discrimination training described above, rats underwent 3 days of training on a RR5 schedule followed by 3 days of training on a RR10 schedule. The following day overexpectation trials were introduced. These sessions contained 12 trials of a single auditory (clicker or white noise; counterbalanced) intermixed with 12 trials of a compound stimulus (e.g., light + the other auditory stimulus; white noise or clicker; see Figure 1). For each trial, whether compound or single, rats could earn a maximum of one pellet if the lever was pressed during stimulus presentation. Rats were assigned to saline (n = 10) or clonidine (n = 11) groups based on response rates in training and received a total of four overexpectation sessions. Infusions of either saline or clonidine (0.6 μg) were given prior to each of these sessions. On the following day, rats were given an extinction test containing four trials of each auditory stimulus (eight trials total) to determine whether previous clonidine treatment impacted the learning that occurred during overexpectation. No infusions were given during testing.

### Histology

After completion of training and testing, rats were overdosed with isoflurane and transcardially perfused with cold, 0.1 M phosphate-buffered saline (PBS) followed by 10% phosphate-buffered formalin. Cryoprotected brains were frozen and sliced using a cryostat (CM1860; Leica, Buffalo Grove, IL, USA), and 40-μm slices mounted, and stained with cresyl violet to verify cannulae placements with the aid of a rat brain atlas [48].

### Statistics

Data are presented as lever-presses per trial and were analyzed with mixed-model analysis of variance (ANOVA). Additional ANOVAs were used to further assess interactions and simple effects where indicated.

## Results

### Histology

Six rats (5 males, 1 female) were excluded from analyses due to off-target cannula placement. Cannula placements for rats included in the behavioural analyses are displayed in Figure 2. Final group sizes were as follows; compound extinction saline (n = 12) and clonidine (n = 9); overexpectation saline (n = 9) and clonidine (n = 9).

**Figure 2.**
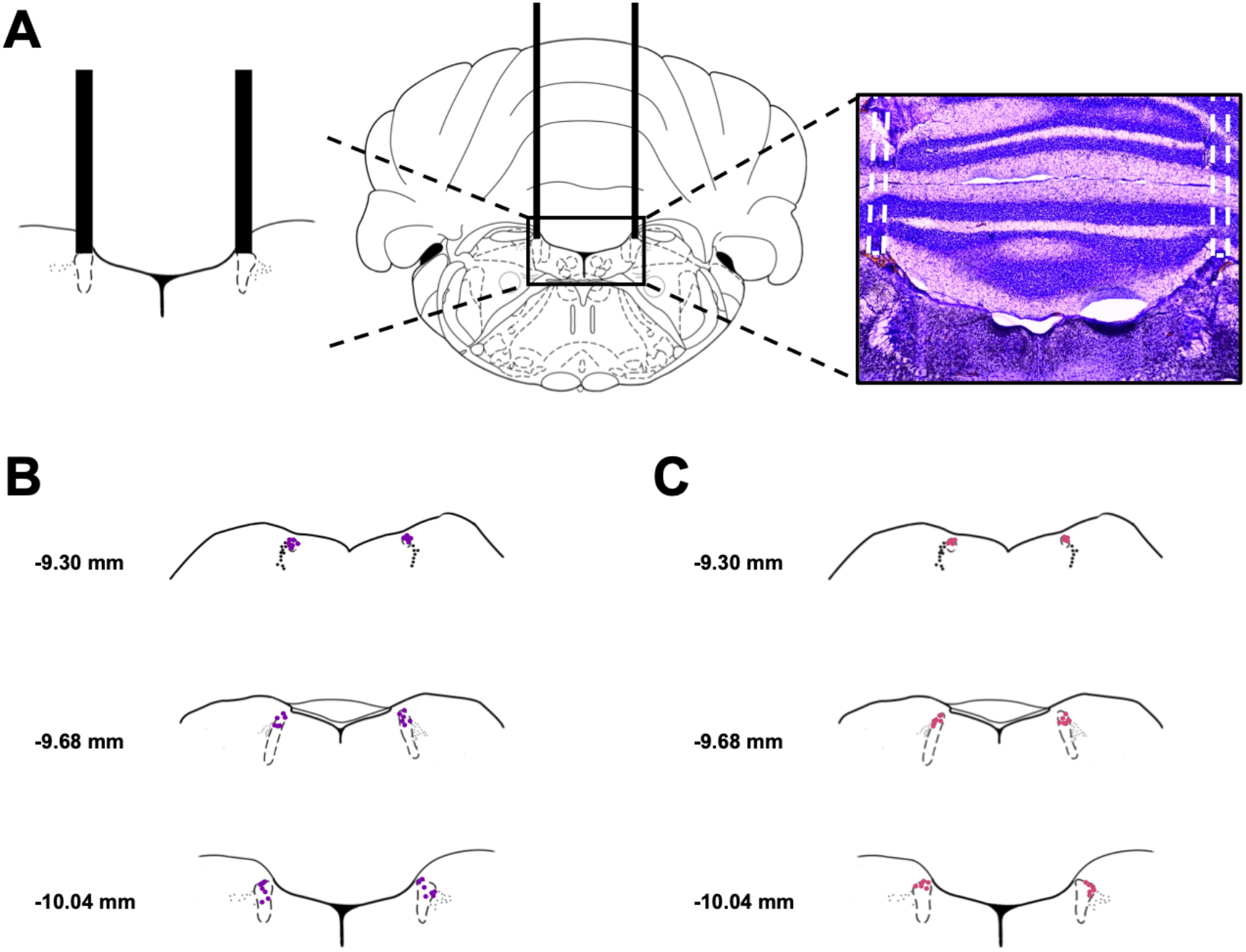
Histology. **(A)** Schematic of the internal cannula target placement (AP: -10.0 mm; ML: ±1.3 mm; DV -7.4 mm) directly above the LC (left) and a representative image of cannula placement after cresyl violet staining (right). **(B)** Diagram of cannula tip placement for all animals included in the compound extinction experiment. **(C)** Diagram of cannula tip placement for all animals included in the overexpectation experiment. Coordinates and images are adapted from Paxinos and Watson 5^th^ Edition [48].

### Compound Extinction

Clonidine and saline groups showed similar performance on the final day of training [Figure 3A; F(1, 18) = 0.034, p = 0.855]. Responding in both groups decreased across the first two days of extinction evidenced by an effect of day [Figure 3B; F(1, 18) = 37.395, p = 0.001], but no effect of group [F(1,18) = 0.015, p = 0.903] and no interaction between these factors [F(1,18) = 0.398, p = 0.536]. Evidence of summation when the compound was introduced was found in the increase in responding from Day 2 of extinction, when stimuli were presented separately, to Day 3 when two stimuli were presented together [effect of day; F(1,18) = 9.084, p = 0.007]. This increase was similar for the two groups as there was no effect of group or day x group interaction [Fs < 1]. There was no difference between groups in responding in the test of spontaneous recovery 4 weeks later [Figure 3C; F(1, 18) = 0.526, p = 0.478]. To confirm spontaneous recovery despite low overall responding, we compared the final two trials of extinction to the first two trials of testing which demonstrated an increase in responding across time [Figure 3D; F(1,18) = 32.763, p = 0.001], but no effect of group [F(1, 18) = 1.687, = 0.210] and no time x group interaction [F(1,18) = 1.033, p = 0.323]. Because spontaneous recovery was weak and transient which may have masked group differences, to further assess retention of extinction, we compared reacquisition of responding in a session where lever-pressing again earned reward. Two animals failed to earn any reward in this session (one from each group) and were excluded from further analyses. The previously saline-treated animals were slower to resume responding consistent with stronger extinction compared to the previously clonidine-treated group that rapidly resumed responding when reward was again available [Figure 2E, F; F(1,16) = 6.808, p = 0.019].

**Figure 3.**
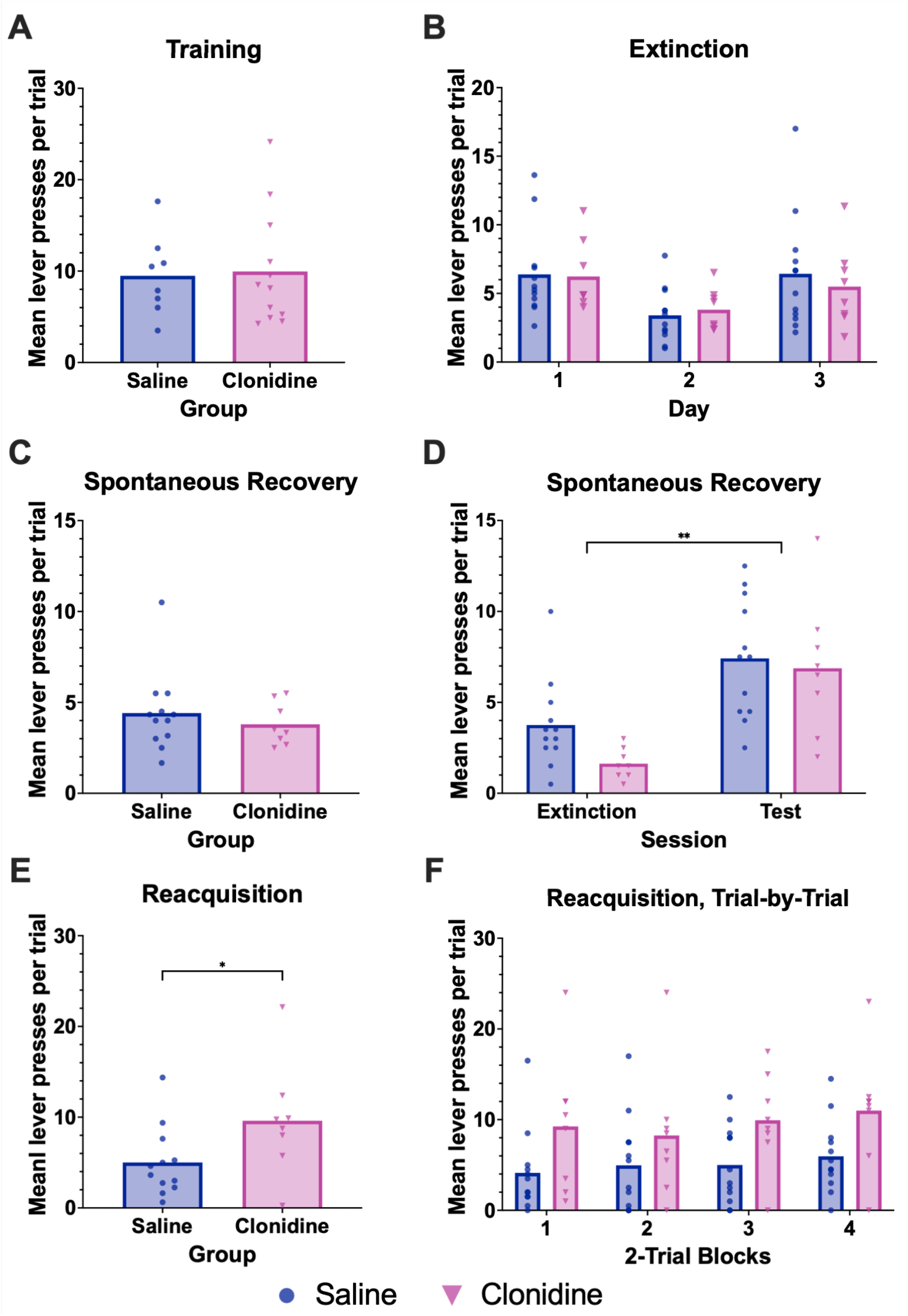
Inhibition of the LC with clonidine attenuates compound extinction allowing rapid reacquisition of responding. **(A)** Responding did not differ between groups on the last day of training, p = 0.855. **(B)** Responding in both groups decreased across the first two days of extinction [p = 0.001]. Further, we observed an increase in responding from Day 2 of extinction, when stimuli were presented separately, to Day 3 when two stimuli were presented together as a compound [p = 0.007], indicating the presence of summation. There was no significant difference between groups or day x group interactions [p’s > 0.05]. There was no significant difference between groups at the spontaneous recovery test conducted 4 weeks after the compound extinction training session (**C**; p = 0.478). **(D)** To confirm spontaneous recovery, we compared the final two trials of extinction to the first two trials of testing and observed an increase in responding across time [p = 0.001] but not across group [p = 0.210]. **(E, F)** When further examining retention of extinction, we found that animals in the saline group were slower to resume responding compared to the clonidine group that rapidly resumed responding when reward was again available [p = 0.019], consistent with weakened retention of extinction learning in the clonidine group. Bars represent group means, with individual data points plotted over top. Clonidine is represented by pink, triangular data point. Saline is represented by blue, circular data points. *, **, *** = p < 0.05, p < 0.01, p < 0.001.

### Overexpectation

On the final day of training were no effects of stimulus [Figure 4A; to-be single, to-be overexpectation; F(1,16) = 0.338, p = 0.542], or group (clonidine, saline; F(1,16) = 0.605, p = 0.448), and no interaction between these factors [F(1,16) = 0.757, p = 0.397]. On the first day of overexpectation, the compound trials produced robust summation. Greater responding was evident in compound (overexpectation) trials relative to single trials [Figure 4B; F(1, 16) = 46.520, p = 0.001]. The clonidine group responded somewhat less overall resulting in a marginal effect of group [F(1,16) = 4.327, p = 0.054] but there was no interaction between stimulus and group indicating elevated responding to the stimulus compound in both groups [F(1,16) = 1.477, p = 0.242].

**Figure 4.**
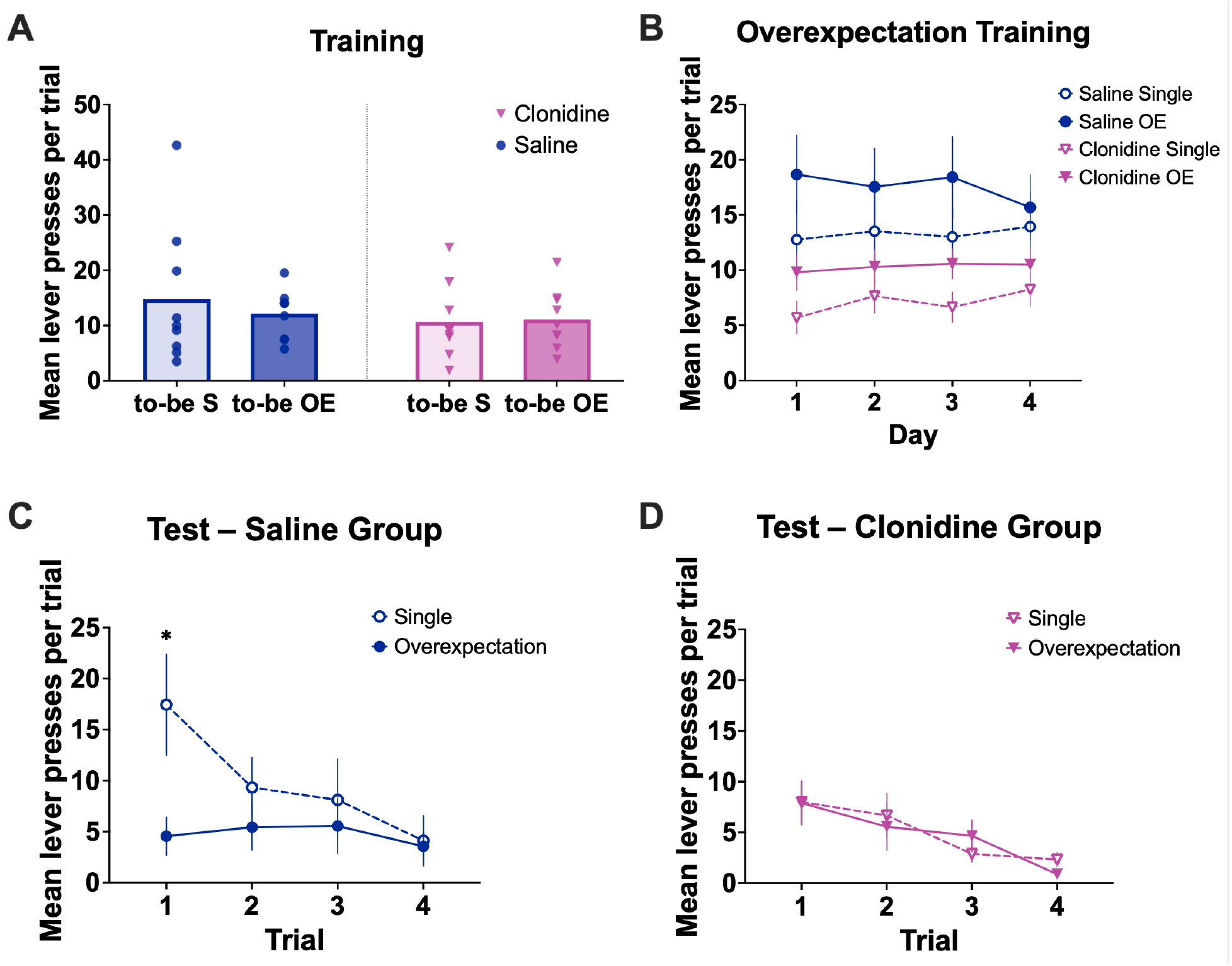
Inhibition of the LC with clonidine attenuates learning driven by a reduction in reward in the overexpectation task. **(A)** On the final day of training, there was no difference between the to-be single (to-be S) and to-be overexpectation (to-be OE) stimuli [p = 0.542], nor the saline and clonidine groups [p = 0.448], and no interaction between these factors [p = 0.397]. **(B)** On the first day of overexpectation training, responding was significantly elevated in compound (overexpectation) trials relative to single trials [p = 0.001]. By the fourth day of overexpectation training, responding to the compound and single stimuli had converged, with no effect of stimulus [p = 0.072], nor group [p = 0.118], and no interaction between these factors [p = 0.809]. **(C)** Early in the test, the stimulus that was previously part of the compound elicited less responding than the stimulus that was always presented alone for the saline group [stimulus x trial interaction; p = 0.037]. Simple effects analyses further confirmed a significant difference between stimuli at trial 1 [p = 0.032] but not at trial 4 [p = 0.757]. This difference was absent in the clonidine group, with no significant difference in responding between stimuli (**D**; p = 0.325). *, **, *** = p < 0.05, p < 0.01, p < 0.001.

Over the course of overexpectation training, responding to the compound and single stimuli converged. By the fourth day of overexpectation, there was no longer an effect of stimulus [Figure 4B; F(1, 16) = 3.717, p = 0.072], nor group [F(1,16)= 2.731, p = 0.118], and no interaction between these factors [F(1,16) = 0.061, p = 0.809].

The data from the test session demonstrate that for the saline group, the stimulus that was previously part of the compound evoked less responding than the stimulus that was always presented alone but that this difference was absent for animals that previously received clonidine [Figures 4C, D]. ANOVA demonstrated no main effect of stimulus [F(1,16) = 2.369, p = 0.143] or stimulus x group interaction [F(1,16) = 1.981, 0.143]. There was however an effect of trial [F(1,16) = 10.522, p= 0.001], a stimulus x trial interaction [F(3,16) = 2.834, p = 0.048] and a stimulus x trial x group interaction [F(3,48) = 3.135, p = 0.034]. To explore the 3-way interaction we examined the effects of stimulus and trial separately for each group. For the saline group, there was no effect of stimulus [F(1,8) = 2.281, p = 0.169], but an effect of trial [F(3, 24)= 7.648, p = 0.001] and a stimulus x trial interaction [F(3,24) = 3.326, p = 0.037]. As suggested by Figure 4C early in the test the stimulus that was previously part of the compound evoked less responding than the stimulus that was always presented alone. Simple effects analyses confirmed an effect of stimulus on trial 1 [F(1,8) = 6.762, p = 0.032] but responding to the two stimuli was equivalent by trial 4 [F(1,8) = 0.103, p = 0.757]. For the clonidine group [Figure 4D], there was an effect of trial, with responding decreasing across trials [F(3,24) = 4.023, p = 0.019], but there was no effect of stimulus [F(1,8) = 0.090, p = 0.772] and no stimulus x trial interaction [F(3,24) = 1.216, p = 0.325] suggesting responding to the two stimuli was equivalent throughout the test session.

## Discussion

These results extend previous findings using systemic pharmacological treatments [15] and optogenetic manipulation of the LC [29,39] and provide further evidence that activity of the LC contributes to the long-term retention of appetitive extinction. Importantly, these data demonstrate that LC activity, and presumed NA release, also contribute to learning that occurs in the overexpectation task, where reward continues to be delivered, but is less than expected based on the predictive stimuli that are present [5,7]. These findings are important because they suggest NA is not uniquely involved in learning about the absence of a reinforcer, but its effects extend to other types of learning driven by a negative prediction error.

From a behavioural perspective, evidence of summation, measured as greater responding to a stimulus compound than to the elements of that compound, supports the idea that animals base their expectations on the combination of predictions from the stimuli available. Importantly, we observed significant summation in the early trials of each task after which responding gradually decreased as predicted by the Rescorla-Wagner model [5] and consistent with the idea that the animals’ prediction for the compound is gradually updated as they gain experience with the new reinforcer (none in extinction, or smaller in overexpectation). While acute clonidine reduced overall responding, greater responding to a stimulus compound was observed in both tasks indicating that clonidine did not impair detection of the new stimulus configurations. However, the updating of stimulus-based predictions when predicted reward was omitted or reduced was impaired when the LC was inhibited resulting in group differences at test potentially due to effects on consolidation [1,49,50].

Clonidine treatment during extinction resulted in faster reacquisition of responding to a previously extinguished stimulus indicating poor long-term expression of extinction and suggesting that the effects of systemic drug administration are likely due to effects on NA released by LC neurons. One difference between the current results and those of previous pharmacological studies is that systemic propranolol administration has been observed to reduce responding to the stimulus compound, which then resulted in greater spontaneous recovery at test. Here, inhibiting the LC did not significantly reduce responding to the compound which could be due to differences in systemic vs. local administration or the specific pharmacological agent (propranolol vs. clonidine). Nonetheless, the presumed reduction in NA release likely affected plasticity at LC targets [18,29] which could account for the effects seen at test. As the LC projects broadly throughout the brain, future studies should examine which of these targets are responsible for the effects of LC inactivation although the amygdala and infralimbic cortex are strong candidates [1,18,29,49,50]. These results may appear at odds with our recent demonstration that optogenetic inhibition of the LC did not impair extinction [39]. However, in that study, the halorhodopsin virus was expressed in only approximately 55% of neurons and the remaining population may have been sufficient to support extinction learning. The clonidine treatment here may have had a more complete effect.

While multiple studies have now implicated NA in extinction [15–18,21,22,29,39], here we explored whether NA is also important for overexpectation. Critically, during testing, saline-treated animals responded less to the overexpectation stimulus consistent with the predicted loss of associative strength across compound trials compared to the single stimulus. This difference was not seen in animals previously treated with clonidine suggesting that inhibition of the LC interfered with the learning that occurred for saline-treated animals during overexpectation training. These results suggest that NA also contributes to learning about a reduced reward, but where reinforcement is still present, thus extending its role beyond extinction. The overexpectation results are also important because this procedure provides an opportunity to reduce the influence of predictive stimuli under circumstances where omitting the outcome entirely may not be possible or practical (e.g., in cases of obesity or eating disorders where the individual must continue to eat). Discovery that LC activity contributes to this effect suggests that enhancing noradrenaline could enhance learning in this paradigm which could have therapeutic benefits. While less is known about the neural circuitry that underlies overexpectation, several studies implicate the central amygdala [51,52]. While the substrates of extinction and overexpectation might be expected to overlap thus pointing to additional candidate regions, they can also be dissociated at the level of the infralimbic cortex [53]. Which regions are important for the effects of NA in this task should be explored in future studies. Further, as there is some evidence that the LC can corelease NA and dopamine [54,55], experiments using sensors specific to each (e.g., GRAB_NE_ or GRAB_DA_) [56,57] potentially alongside experiments that inhibit NA or dopamine receptors could isolate the contribution of each neuromodulator to learning in extinction and overexpectation paradigms.

Finally, studies that directly measure the LC response when a prediction error is introduced would give insight into how the endogenous NA system is engaged by prediction errors. Further, while here, we find evidence that the LC is important for two types of learning driven by negative prediction error, future studies should examine whether the LC has a general role in prediction error and is also involved in learning where that error is positive (i.e. reward is greater than expected) or whether it’s role is specific to negative error, potentially complementing the established role of dopamine in signaling positive prediction error [58–60].

The ability to predict significant events is highly adaptive. It is also essential to be able to update previously learned associations when conditions change. The data here demonstrate that activity of the LC is important for learning to reduce responding in both extinction and overexpectation paradigms. Understanding how the brain updates previously learning to reduce the acquired power of predictive stimuli is important for developing more effective therapies that aim to reduce the influence of such stimuli and to do so with more lasting effects.

## Data Availability Statement

Raw data is publicly available through the University of Toronto’s data repository: https://doi.org/10.5683/SP3/VDGTGO

## Author Contributions

All authors contributed extensively to the work presented in this paper. SKC and LHC designed the experiments. SKC and AKB performed the experiments. All authors contributed to the preparation of the manuscript.

## Supplemental Materials

### Dose Response Testing

To establish an effective dose of clonidine that did not produce sedative effects, we examined the effects of clonidine on reinforced lever-pressing. Rats (n = 8) were trained to lever press for a food pellet reward during stimulus presentations on a continuous reinforcement schedule. The effects of a LC-targeted infusion of 0.2 μL of sterile saline, or 0.6, 1.2, or 2.4 μg of the alpha-2 adrenergic receptor agonist, clonidine (clonidine hydrochloride; MW: 266.55 g/mol; Sigma-Aldrich) dissolved in 0.2 μL of sterile saline on reinforced lever-pressing were assessed within subjects across days according to a Latin square design. A day without infusions was given between doses to avoid any cumulative drug effects. Based on these results, we proceeded with a dose of 0.6 μg of clonidine for the following tests (see Figure S1).

**Supplemental Figure 1.**
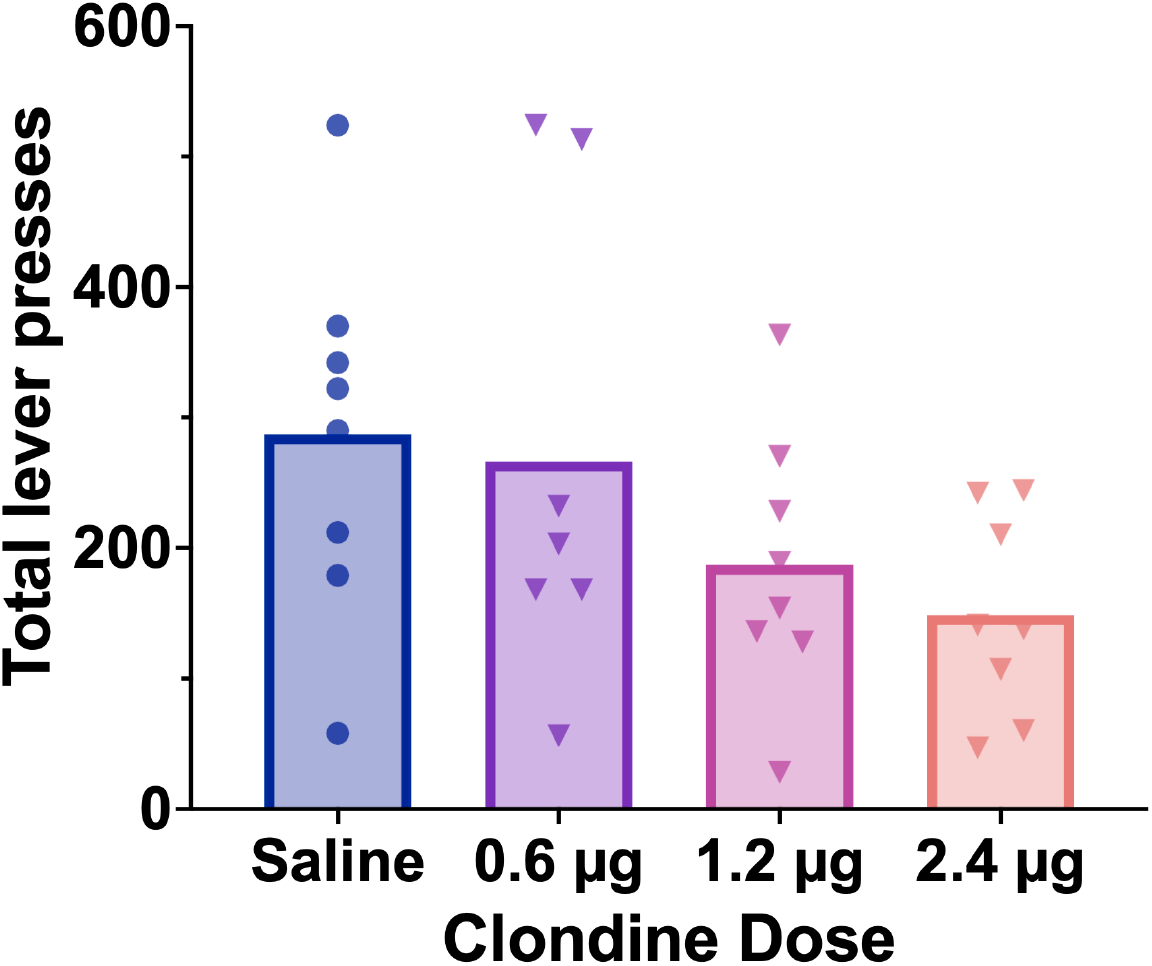
An 0.6 μg dose of clonidine in the LC does not reduce overall responding. Before conducting our primary experiments, we sought to identify an appropriate dose of clonidine that suppresses noradrenergic function in the LC but was not generally deleterious to food reward-driven behaviours. We examined the effects of an LC-targeted infusion of the alpha-2 adrenergic receptor agonist, clonidine (sterile saline, 0.6, 1.2, or 2.4 μg) on food reward-driven lever pressing. We identified 0.6 μg as the highest dose that did not produce suppression of lever pressing behaviours. Bars represent group means, with individual data points plotted over top.

